# The Kocurious case of Noodlococcus: genomic insights into *Kocuria rhizophila* from characterisation of a laboratory contaminant

**DOI:** 10.1101/2025.05.28.656266

**Authors:** Gregory E. McCallum, Siu Fung Stanley Ho, Elizabeth A. Cummins, Alex J. Wildsmith, Ross S. McInnes, Christoph Weigel, Lok Yee Sylvia Tong, Joshua Quick, Willem van Schaik, Robert A. Moran

## Abstract

The laboratory contaminant Noodlococcus was named for its coccoid cells and unusual colony morphology, which resembled a pile of noodles. Along with laboratory characterisation and electron microscopy, we generated a complete Noodlococcus genome sequence using Illumina and Oxford Nanopore data. The genome consisted of a single, circular, 2732108 bp chromosome that shared 97.5% Average Nucleotide Identity (ANI) with the *Kocuria rhizophila* type strain TA68. We identified genomic features involved in replication (*oriC*), carotenoid synthesis (*crt*), and genome defence (CRISPR-Cas), and discovered four novel mobile elements (IS*Krh4-7*). Despite its environmental ubiquity and relevance to food production, bioremediation, and human medicine, there have been few genomic studies of the *Kocuria* genus. We conducted a comparative, phylogenetic, and pangenomic examination of all 257 publicly available *Kocuria* genomes, with a particular focus on the 56 that were identified as *K. rhizophila*. We found that there are two phylogenetically distinct clades of *K. rhizophila*, with within-clade ANI values of 96.7-100.0% and between-clade values of 89.5-90.4%. The second clade, which we refer to as *K. pseudorhizophila*, exhibited ANI values of <95% relative to TA68 and constitutes a separate species. Delineation of the two clades would be consistent with the rest of the genus, where all other species satisfy the 95% ANI threshold criteria. Differences in the *K. rhizophila* and *K. pseudorhizophila* pangenomes likely reflect phenotypic as well as evolutionary divergence. This distinction is relevant to clinical and industrial settings, as strains and genomes from both clades are currently used interchangeably, which may lead to reproducibility issues and phenotype-genotype discordance. Investigating an innocuous laboratory contaminant has therefore provided useful insights into the understudied species *K. rhizophila*, prompting an unexpected reassessment of its taxonomy.

**Impact statement:** Bacterial genome sequence databases are dominated by a relatively small number of medically relevant genera, while most of the global bacterial population’s diversity is largely uncharacterised. *Kocuria* is a widespread bacterial genus with industrial and medical relevance. Despite its ubiquity, only 22 complete and 235 draft *Kocuria* genomes were publicly available at the outset of this study. Our phylogenetic and pangenomic examination of all available *Kocuria* genomes was the first for this genus, providing insights into its diversity and taxonomy. Most notably, we found that *Kocuria rhizophila* is comprised of two clades that are sufficiently divergent to constitute different species, but are frequently used interchangeably in experimental and genomic research. The complete, high-quality Noodlococcus genome generated and characterised here can serve as a reference for true *K. rhizophila*, particularly while there is only a draft genome sequence available for type strain TA68.

**Data summary:** Sequencing reads and the assembled Noodlococcus genome are available from NBCI BioProject accession PRJNA835814 and BioSample accession SAMN28111796. The complete sequence of the Noodlococcus chromosome can be found in the GenBank nucleotide database under accession number CP097204.1. Entries for the novel insertion sequences IS*Krh4* to IS*Krh7* can be found in the ISFinder database (https://isfinder.biotoul.fr).

## Introduction

*Kocuria* is a bacterial genus in the family Micrococcaceae that was first distinguished from *Micrococcus* in 1995 [1]. As of September 2024, there were 27 *Kocuria* species listed in the National Centre for Biotechnology Information (NCBI) Taxonomy Browser, 22 of which were represented by genome sequences. All *Kocuria* are Gram-positive cocci that arrange in mixtures of short chains and disordered clusters, forming colonies that are usually pigmented [1, 2], with yellow [3], orange [4], and pink [5] colonies described. *Kocuria* species have been isolated from diverse terrestrial and marine environments, and strains have exhibited a wide array of phenotypes, including tolerance of cold temperatures, radiation, pH extremes, or salinity [4, 6–10].

*Kocuria* has been found as a contaminant in clinical and research laboratories [11], likely because of its environmental ubiquity and presence on human skin. Mistaking *Kocuria* in clinical samples for contaminants can lead to misdiagnoses in rare cases where *Kocuria* causes human infections [12]. Reported *Kocuria* infections have included cases of peritonitis [13, 14], bacteraemia [15–18], meningitis [19], endocarditis [20–22], postsurgical infections [23], and infections of various other body sites [2, 24]. Infections are typically responsive to antimicrobial chemotherapy as *Kocuria* are generally susceptible to antibiotics, apart from their intrinsic resistance to nitrofurantoin and furazolidone [2, 3, 5].

*Kocuria rhizophila* was named in 1999 following characterisation of the type strain TA68, which was isolated in 1995 from the rhizosphere of *Typha angustifolia* (narrowleaf cattail) growing on a floating mat in the Soroksár tributary of the Danube river, Hungary [3]. *K. rhizophila* has since been isolated from a wide range of terrestrial and aquatic environments such as Siberian permafrost [25], peat soil [26], waterfalls [27], and deep-sea sponges [28]. It has been found on food products [29], and in close association with humans [30], plants [3, 7, 31, 32], and animals [33–35]. Strains of *K. rhizophila* have been used routinely as controls for antibiotic sensitivity testing [36].

Despite its ubiquity and relevance to human health, there is a sparsity of *Kocuria* genome data in public databases. The first complete *Kocuria* genome was the soil-derived *K. rhizophila* strain DC2201, which was published in 2008 [37]. As of September 2024, there were 257 *Kocuria* genomes in the NCBI database, 22 of which were complete (Table S1). Complete genome sequences have revealed that *K. rhizophila* has one of the smallest actinobacterial chromosomes, ranging from 2.6-2.8 million base pairs [33, 37]. Given their vast potential for diverse metabolite production, the need for expanded knowledge of actinobacterial genomics has been expressed recently [38]. While the number of publicly available *Kocuria* genomes has been increasing over recent years, we are aware of only a single study that has performed comparative genomic analysis of six *Kocuria* genome sequences from four species [39].

In August 2019 we found an unusual colony growing on an agar plate that had been left on a laboratory bench for 11 days. The colony was raised, yellow-pigmented, and looked like a pile of noodles. After a Gram stain revealed that it was a Gram-positive coccus, we named the isolate Noodlococcus and sought to identify it. Generating a complete Noodlococcus genome sequence revealed that it was a strain of *Kocuria rhizophila*. We characterised the Noodlococcus genome and used it as a vehicle for comparative analyses of *Kocuria*, focusing particularly on the phylogeny and pangenome of *K. rhizophila*. Our investigation uncovered previously unrecognised diversity in *K. rhizophila*, with consequences for the classification of widely used type strains and genome sequences.

## Materials and Methods

### Media, isolation, and culture conditions

The original Noodlococcus colony was found on a brain heart infusion (BHI) agar plate that had been used to culture *Enterococcus faecium* in a laboratory at the University of Birmingham, United Kingdom. An overnight *E. faecium* culture in BHI broth had been diluted and spread on the BHI plate to obtain single colonies. After overnight incubation at 37°C, *E. faecium* colonies were picked using sterile pipette tips and transferred to new agar plates. The plate that colonies had been picked from was left in ambient conditions on a laboratory bench for 11 days before Noodlococcus and re- grown *E. faecium* colonies were observed. Noodlococcus was subsequently re-streaked onto a BHI agar plate and incubated at 37°C overnight. A single colony was then picked and grown in BHI broth at 37°C with shaking at 200 revolutions per minute (rpm). Gram staining was performed as described by Coico [40].

A variety of conditions were tested to determine the optimal growth conditions for Noodlococcus. Growth in tryptic soy broth (TSB), BHI broth, lysogeny broth (LB), and nutrient broth was compared by adding 5 μL of overnight culture to 100 mL of broth in 500 mL conical flasks. The flasks were then incubated for 24 hours at 30°C with shaking at 200 rpm. Aggregated Noodlococcus cells were broken up by pipetting before the optical density at 600 nm (OD_600_) was measured from 1 mL of culture. Three technical replicates were used for each condition.

To determine the optimal temperature for growth, culture turbidity was measured at 4, 10, 15, 20, 25, 28, 30, 37, 40, 45, and 50°C. A single Noodlococcus colony was picked and added to 50 μL TSB. After mixing by pipetting, 5 μL of this inoculum was added to 6 mL TSB in a 30 μL universal container for each temperature condition (in biological triplicate, each with 3 technical replicates). OD_600_ measurements were taken for 1 mL of the samples to establish a baseline reading. Samples were incubated with shaking at 200 rpm. After 24 hours, cultures were mixed by pipetting to break up cell aggregates, and OD_600_ was measured again. Growth in a range of NaCl concentrations (0, 1, 2, 3, 5, 7, 10, 15, 20% weight/volume) and pH conditions (pH 3, 4, 5, 6, 7, 8, 9, 10, 11, 12) was determined in the same way in TSB at 30°C. For salinity and pH tests, OD_600_ was also measured after 96 hours of growth. To evaluate growth in anaerobic conditions, Noodlococcus was streaked onto TSB agar plates and incubated for 96 hours in a MACS MG-500 Anaerobic Chamber Workstation (Don Whitley) set to 37°C and connected to a 5% Carbon Dioxide, 5% Hydrogen/Nitrogen (Anaerobic) Cylinder (BOC). For all subsequent experiments, Noodlococcus was cultured in TSB at 30°C with 200 rpm shaking, and on TSB agar plates at 30°C, unless stated otherwise.

### Electron microscopy

Overnight Noodlococcus culture was fixed in 2.5% glutaraldehyde for 30 minutes at 4°C. Scanning electron microscopy (SEM) was conducted with a SEM-Zeiss EVO15 VP ESEM microscope at the Centre for Electron Microscopy at the University of Birmingham.

### Biochemical analysis

Noodlococcus was sent to DSMZ (Braunschweig, Germany) for fatty acid composition analysis. An oxidase test was carried out by smearing a colony onto an oxidase test strip (Merck) and observing for a change in colour after 10 seconds. *Pseudomonas aeruginosa* PAO1 was used as a positive control, and *Escherichia coli* DH5α as a negative control. A catalase test was performed by smearing a colony onto sterile glass, adding one drop of 3% hydrogen peroxide, and observing for the production of bubbles. *E. coli* DH5α was used as a positive control, and *E. faecium* 64/3 as a negative control.

### Antibiotic susceptibility testing

The susceptibility of Noodlococcus to ampicillin, cefotaxime, ceftazidime, ciprofloxacin, colistin, erythromycin, nitrofurantoin, and tetracycline was measured using the broth microdilution method [41] and interpreted against the EUCAST breakpoints [42]. Assays were performed in biological triplicate and the mode of the three replicates was recorded.

### DNA extraction

1 mL of overnight culture was homogenised by vigorous vortexing for one minute and DNA was extracted using the Wizard Genomic DNA Purification Kit (Promega) using their Gram-positive protocol with the inclusion of lysozyme (10 mg/mL; Sigma-Aldrich). DNA concentrations were quantified using the Qubit dsDNA BR assay kit (Thermo Fisher). DNA quality was assessed using a NanoDrop 2000 spectrophotometer.

### Genome sequencing and assembly

Short read DNA libraries were prepared with the Nextera XT library prep kit (Illumina). Shotgun sequencing was carried out by MicrobesNG (Birmingham, United Kingdom) using the HiSeq 2500 sequencing platform (Illumina) which generated 150 bp paired-end reads. Fastp v0.23.2 [43] was used to trim adapter sequences, and to remove both low quality and duplicate reads (--dedup).

Long-read sequencing libraries were constructed with the Ligation Sequencing Kit SQK-LSK109 (Oxford Nanopore Technologies (ONT)) with the following minor adjustments. An additional Ampure bead clean-up was carried out before DNA repair and end prep to improve ligation efficiency. To further increase ligation efficiency, incubation times for end repair, dA-tailing, and ligation were increased to 30 minutes [44]. Long-read libraries were sequenced on a GridION (ONT) using a FLO-MIN106D R9.4.1 flow cell (ONT), MinKNOW 5.0.5 (ONT), and a 72-hour run script with active channel selection enabled. The sequencing signal was basecalled using Guppy v6.0.1 (ONT) super accuracy mode (--chunk-size 3000). The bottom 5% of reads by quality score and reads less than 1 kilobase (kb) were removed with Filtlong v0.2.1 [45].

Long-read assembly was performed using the consensus assembler Trycycler v0.5.3 [46]. Briefly, reads were subset into 12 different samples which were fed into three assemblers: Flye v2.9 [47], Miniasm+Minipolish v0.3, v0.1.3 [48, 49], and Raven v1.6.0 [50] (4 samples each). Contigs from all three assemblies were clustered, reconciled, aligned, and partitioned to generate a consensus assembly. The accuracy of the consensus assembly was increased with Medaka v1.0.6 [51]. Finally, the assembly was polished with the quality-controlled Illumina reads using PolyPolish v0.5 [52] and then POLCA (MaSuRCA suite v4.0.9) [53]. The genome was annotated using Bakta v1.9.4 [54].

### Genome feature identification and annotation

CRISPR repeats were identified manually in the region downstream of Cas genes. Replication origin prediction was performed as outlined previously [55]. Insertion sequences were found by sequentially comparing 100 kb segments of the Noodlococcus chromosome to its entire sequence in order to identify regions of >500 bp that occurred at multiple positions and shared >99% nucleotide identity. These regions were checked for the presence of transposase genes, and putative transposase sequences were used to query the ISFinder database [56]. IS were assigned to families based on transposase identities, with inverted repeats and target site duplications identified manually when relevant. The complete genome of *K. rhizophila* 28R2A-20 (GenBank accession CP072262 [28]) was used to identify naïve insertion positions that did not contain IS.

### Phylogenetic analysis

All published *Kocuria* genomes (*n=*257) were downloaded using the NCBI Datasets platform on 21^st^ September 2024. Genome completeness and contamination were assessed using CheckM2 [57]. Genomes with <90% completeness and/or >5% contamination were filtered out. Average Nucleotide Identities (ANIs) were determined by FastANI v1.33 [58]. Each genome was annotated using Bakta v1.9.4 [54]. Panaroo v1.5.0 [59] (--clean-mode moderate) was used to generate a core-genome alignment of the genomes, including the Noodlococcus genome, and IQ-Tree v2.3.6 [60] was then used to infer a maximum likelihood phylogenetic tree using the best fit nucleotide substitution model (GTR+F+I+R5 for core genome phylogenies) as determined by ModelFinder [61], and 1000 ultrafast bootstrap replicates [62]. A core genome alignment was generated (--clean-mode set to strict) with all the genomes that were monophyletic to the *K. rhizophila* strain TA68 or NCTC8340. Two strains of *K. tytonicola* (strains 473 and DSM 104133) were included in this alignment for use as an outgroup. Representative full length 16S ribosomal RNA (rRNA) genes, annotated by Bakta, were extracted from the genomes that passed CheckM2 quality control (QC) and aligned with MAFFT’s [63] G-INS-i algorithm before a phylogeny was inferred as described above. Genomes containing only partial 16S rRNA genes were removed from this analysis. Phylogenetic trees were visualised using TreeViewer v2.2.0 [64]. A heatmap of the ANI results was generated using the pheatmap R package v1.0.12 [65].

### ANI comparisons

To compare intra- and inter-species ANI values, all genomes that passed CheckM2 QC (*n=*230) were first classified using Genome Taxonomy Database (GTDB) and associated taxonomic classification toolkit (GTDB-Tk) v2.3.2 [66] classify workflow using release 214 of the GTDB-Tk reference package [67]. GTDB-Tk adds alphabetic suffixes to the end of genus or species names if the classification is ambiguous, however for the sake of grouping the *Kocuria* genomes into species for ANI comparisons, these suffixes were removed. Genomes classified outside of the *Kocuria* genus (*n=*13) and genomes only classified as *Kocuria* to the genus level (*n=*16) were discarded. Any *Kocuria* species made up of less than 3 genomes (*n=*6 species out of 16: *K. coralli*, *K. dechangensis*, *K. polaris*, *K. soli*, *K. tytonicola*, and *K. tytonis*) were also discarded (*n=*9 genomes), leaving a total of *n=*192 genomes classified as a *Kocuria* species with >2 other genomes sharing the same species classification. ANIs for these genomes calculated using FastANI were used to make intra- and inter- species comparisons.

### Pangenome analysis

A pangenome was generated from the 51 *K. rhizophila* genomes using Panaroo (-- clean-mode strict) with a 98% sequence identity threshold. Panaroo’s default definitions of core (99 ≤ x ≤ 100%), soft core (95 ≤ x ≤ 99%), shell (15 ≤ x ≤ 95%), and cloud (0 ≤ x ≤ 15%) were used. The twilight analysis package [68] was used to perform population structure-aware gene classification between the two clades. Functional annotation was performed by eggNOG-mapper v2.0 [69] with the eggNOG v5 database [70]. Prophages were detected using geNomad v1.11.0 [71], with quality and completeness determined using CheckV v1.0.3 [72].

## Results

### Morphological characteristics of Noodlococcus

The original Noodlococcus colony was round, raised, and sulphur yellow. It measured approximately 9 mm across and 2 mm tall, with a complex secondary structure that resembled a pile of noodles (Fig. 1a, b). Subsequent streaking on BHI agar produced small (1 mm) yellow colonies after incubation at room temperature or 37°C for 24 hours, which developed a raised central ring structure when left at room temperature for seven days (Fig. S1), and took between two and three weeks to form secondary structures that resembled the original colony. In BHI broth incubated shaking (220 rpm) at 37°C overnight, Noodlococcus produced a single non-diffuse colony-like structure.

**Fig. 1.**
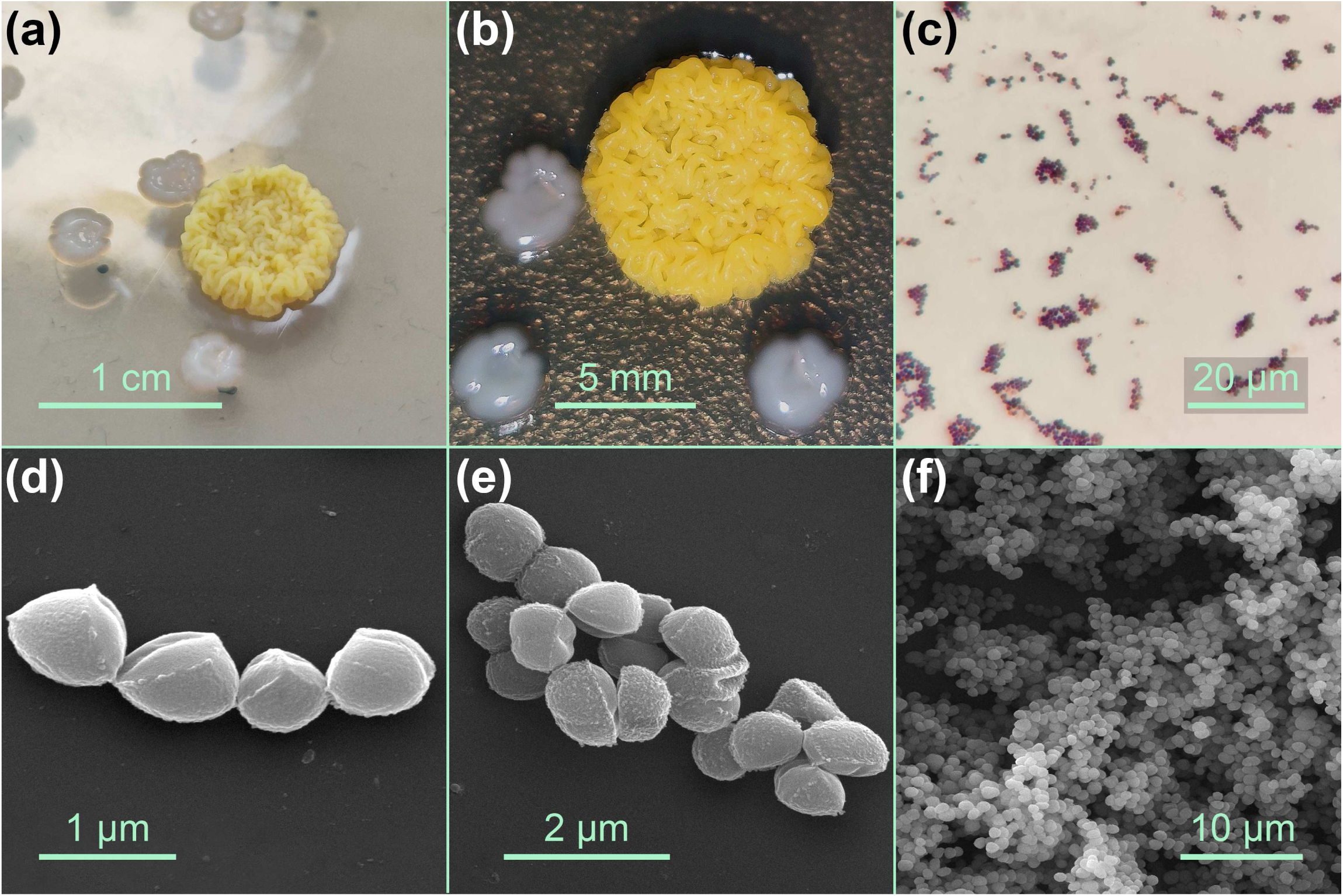
Morphology of laboratory contaminant Noodlococcus. **(a, b)** Photographs of the original Noodlococcus colony that was found growing on brain heart infusion agar. The smaller, white colonies were *Enterococcus faecium* that had re-grown after being picked for further culturing 11 days prior. **(c)** Phase-contrast micrograph of Gram-stained Noodlococcus. **(d-f)** Scanning electron micrographs of Noodlococcus cells.

Gram staining revealed that Noodlococcus was a Gram-positive coccus arranged in short chains or irregular clusters (Fig. 1c). Scanning electron microscopy showed that individual cocci were non-uniform, generally ovoid, and approximately 0.7-1.0 µm long and 0.5-0.8 µm wide (Fig. 1d, e, f).

### Growth conditions and biochemistry

Of the liquid media assessed, Noodlococcus grew optimally in TSB under aerobic conditions (Fig. S2a). It did not grow anaerobically. Noodlococcus grew at temperatures between 20-40°C, with optimal growth at 28°C (Fig. S2b). It grew between pH6 and pH11, with optimum growth at pH7 (Fig. S2c). Growth was reduced in media containing >3% NaCl (Fig. S2d).

Noodlococcus was oxidase-negative and catalase-positive. Fatty acid composition analysis revealed that the major fatty acids present in the Noodlococcus cell wall were anteiso-C_15:0_ (45.7%), anteiso-C_17:0_ (16.9%), and iso-C_15:0_ (14.9%), with closest matches identified in the *Micrococcus-luteus-GC* subgroup C (sim index score 0.521). Noodlococcus was sensitive to penicillin, cephalosporin, fluoroquinolone, macrolide, and tetracycline antibiotics, but resistant to colistin and nitrofurantoin, with MICs of 32 and >256 µg/mL, respectively (Table S2).

### Complete genome of *Kocuria rhizophila* Noodlococcus

The complete Noodlococcus genome was assembled from a combination of Illumina and Nanopore reads using Trycycler. The genome consisted of a single, circular 2,732,108 bp chromosome with an overall G+C content of 70.6%. The Noodlococcus 16S rRNA gene was 99.9% identical to that of the *K. rhizophila* type strain TA68, with 96% query coverage. Confirming its species assignment, we found that the ANI of the Noodlococcus genome relative to that of TA68 was 97.5%. Bakta annotation identified 2,343 open reading frames (ORFs), 46 tRNAs, and 9 rRNAs in the Noodlococcus chromosome. A putative origin-of-replication (*oriC*) was identified between the *dnaA* and *dnaN* genes (positions 601,211-601,811 of GenBank accession CP097204). It includes a DNA unwinding element, DnaA-trio motifs, and two arrays of DnaA boxes, resembling previously characterised actinobacterial chromosomal replication origins [55, 73] (Fig. 2a).

**Fig. 2.**
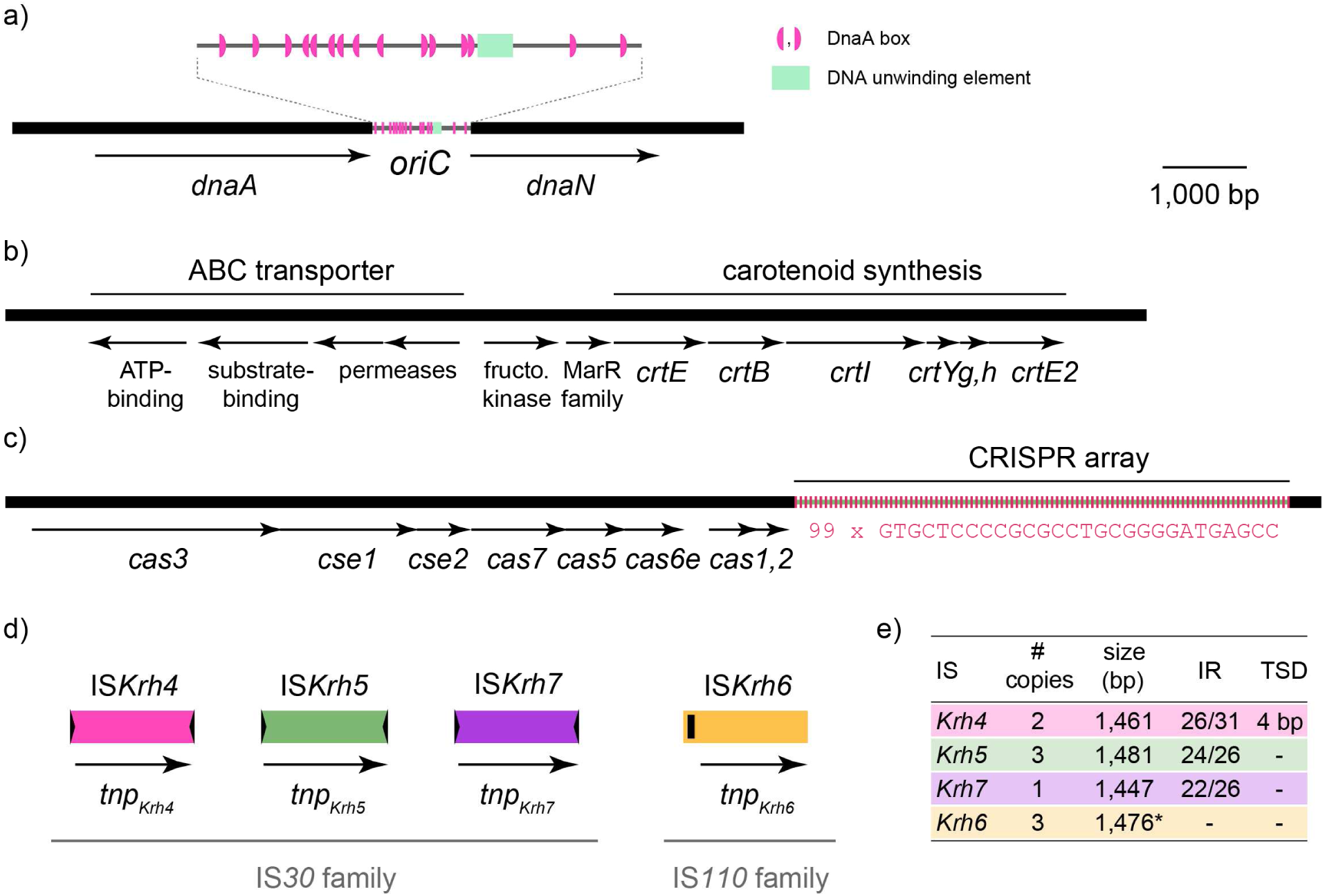
Genomic features of Noodlococcus. All parts drawn to the same scale from GenBank accession CP097204. The extents and orientations of genes are indicated by labelled arrows beneath the horizontal lines that represent segments of the Noodlococcus genome. **(a)** Origin-of-replication. The *oriC* region is magnified 4.5x above to display fine-scale features as indicated in the key to the right. **(b)** Carotenoid synthesis region. The extents of regions that determine carotenoid synthesis and a putative ABC transporter are marked by labelled lines above. **(c)** CRISPR-Cas locus. The extent of the CRISPR locus is marked above. Each short, vertical pink line represents a copy of the 28 bp repeat unit, the sequence of which is shown below. **(d)** Insertion sequences (IS). Coloured vertical lines represent novel IS found in the Noodlococcus chromosome, grouped according to family membership. Terminal inverted repeats are shown as short black arrows and a seekRNA determining region as a small black rectangle. **(e)** IS characteristics. Table outlining the features of novel IS characterised here. * = size estimate that requires experimental validation (see text), IR = terminal inverted repeats (identical bases/total bases), TSD = target site duplication.

As its colour was such a distinctive aspect of its colony morphology, we searched the Noodlococcus genome for pigment determinants. Carotenoids are naturally occurring pigments produced by a wide range of organisms for a variety of purposes [74]. We found a putative carotenoid synthesis cluster that contained six *crt* genes and resembled the gene cluster of *Micrococcus luteus* NCTC 2655 that directs synthesis of the γ-cyclic C_50_ carotenoid sarcinaxanthin, which has been experimentally characterised [75]. The Noodlococcus cluster contains *crtE*, *B*, *I*, *Yg*, *Yh* and *E2* genes (Fig. 2b) that encode proteins with amino acid identities that range from 48.1-75.6% identical to their equivalents from NCTC 2655, but does not contain a gene equivalent to *crtX*. Genes for a MarR-family regulator and fructosamine kinase lie immediately upstream of the *crt* genes, separating them from a set of four genes for a putative ABC transporter (Fig. 2b).

A CRISPR locus was identified downstream of determinants for a type I-E CRISPR-Cas system (Fig. 2c). The locus contained 98 spacer sequences of 33 or 34 bp, interspersed with 99 copies of a 28 bp repeat unit. The spacer sequences were used to query the GenBank non-redundant nucleotide database, and 64/98 returned matches to non-*Kocuria* sequences (Table S3). Fifty-six spacers matched bacteriophage genomes derived from urban environment metagenomic datasets with identities ranging from 90.9-100%, and eight matched chromosomal sequences from various bacterial genera with identities 84.8-93.9%. All 15 of the bacteriophage genomes matched by Noodlococcus spacers were *Caudoviricetes* and have previously been predicted to be lytic [76]. To account for lysogenic bacteriophage, the Noodlococcus chromosome was screened for the presence of prophage regions, but none were found.

Three novel insertion sequences (IS) were identified based on their presence at multiple positions in the Noodlococcus chromosome (Fig. 2d, e). They included two IS*30* family elements, IS*Krh4* (at two chromosomal positions, flanked by distinct 4 bp target site duplications [TSDs]) and IS*Krh5* (at three positions, no TSDs). Another novel element, IS*Krh7*, present at a single chromosomal position, was identified by homology to IS*Krh5*, with which it shared 74.2% nucleotide identity (1,044/1,047 bp). The third IS found at multiple positions was IS*Krh6*, which belongs to the IS*110* family. It has recently been demonstrated that the non-coding region upstream of the transposase gene in IS*110* family elements determines a seekRNA that directs transposition [77]. The size of IS*Krh6* was estimated as 1,476 bp based on the extent of conserved sequence at all three positions in Noodlococcus, which might include the element’s target site (<10 bp), but this would need to be distinguished from the ends of the element experimentally.

### Diversity of publicly available *Kocuria* genomes

A total of 257 *Kocuria* genomes were retrieved from the NCBI database (last search 21^st^ September 2024). These included representatives of 24 *Kocuria* species, with 80 genomes unclassified. A total of 230 (89.5%) passed CheckM2 QC (≥90% completeness, ≤5% contamination). A genus-wide core genome phylogeny revealed a structure that was largely concordant with existing species definitions (Fig. S3). The phylogeny was supplemented by using FastANI to compare ANI across all genomes. ANI is used widely for assessing species boundaries, with >95% ANI a typical threshold for genomes belonging to the same species [58]. Most *Kocuria* species definitions were supported by ANI values (Fig. S3). Apart from exceptions detailed below, inter-species ANI values ranged from 77.7-88.8%, while intra-species ANI values ranged from 94.5-100%. Two genomes were outliers — *Kocuria palustris* DE0549 and *Kocuria rosea* TA28 — which had ANI values of 85.9% and 86.0% to their respective type strains (Fig. S3, Fig. S4). These were likely mislabelled genomes that either represented novel *Kocuria* species or were a result of assembly chimeras, but comparison to further closely-related genome sequences would be needed to confirm this. A major exception to the largely concordant data was *K. rhizophila*, which included Noodlococcus and accounted for 22.2% of *Kocuria* genomes examined here.

Fifty-six genomes were labelled as *K. rhizophila* with a wide range of isolation sources including human, animals or animal products, and diverse environments (Table 1). Six of these genomes did not pass QC, including four which were flagged or suppressed by GenBank due to contamination, inappropriate genome size, or other genome content issues (Table S4). In the core genome phylogeny, all genomes labelled *K. rhizophila* clustered in two monophyletic clades except for L3_129_000G1_dasL3_129_000G1_metabat.metabat.108 and TNDT1 which were located within the *K. salsicia* and *K. flava* clusters (Fig. S3). These genomes also had whole genome ANIs of 85.7% and 80.7% relative to *K. rhizophila* type strain TA68, respectively, providing further evidence that they have been mislabelled. Three genomes (HMSC066H03, BT304, and APC 4018), that had been labelled as unclassified *Kocuria* sp. in GenBank, also clustered within the two clades (Fig. S3), leaving a total of 51 *K. rhizophila* genomes (Table 1).

**Table 1.**
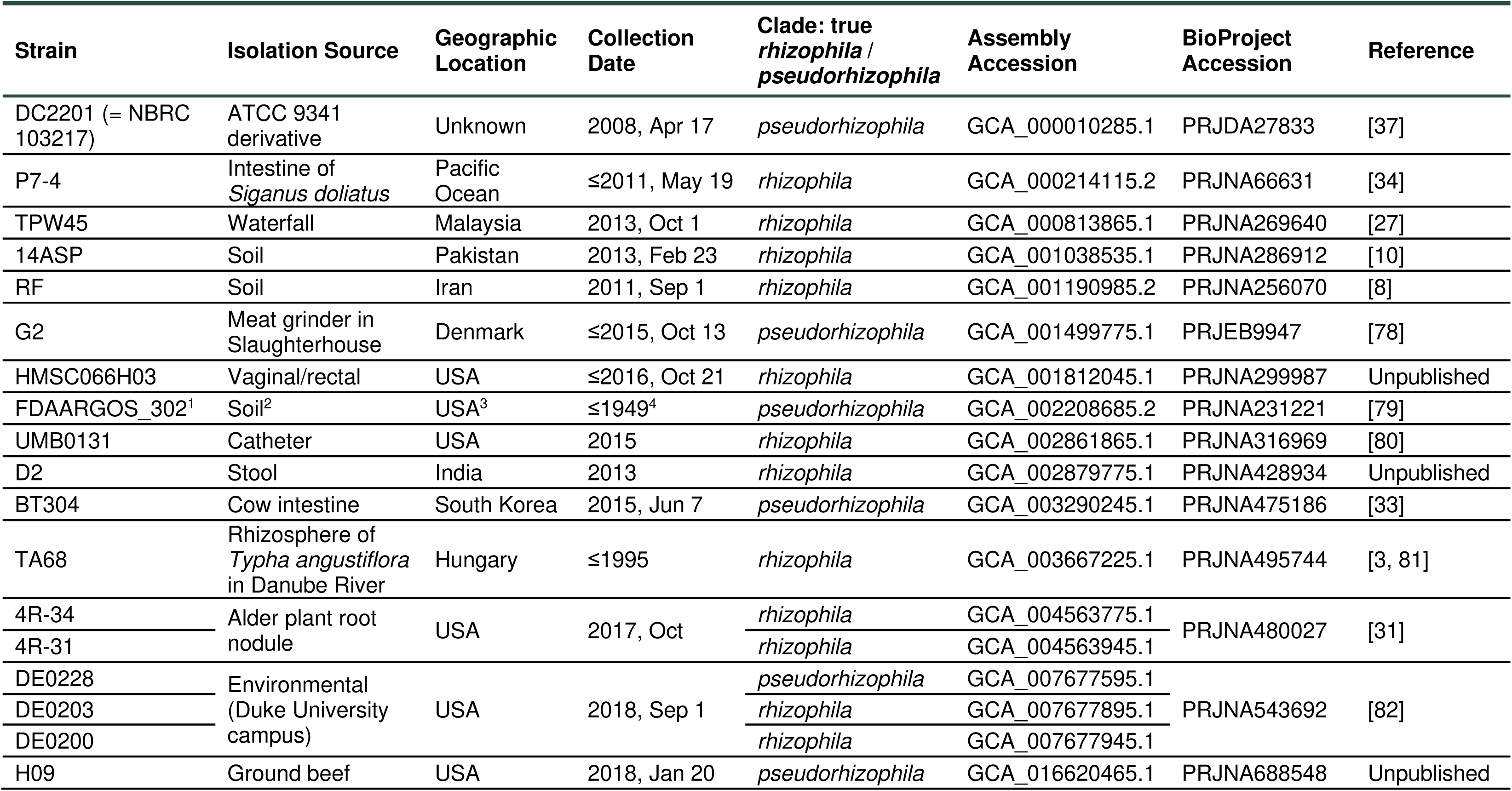
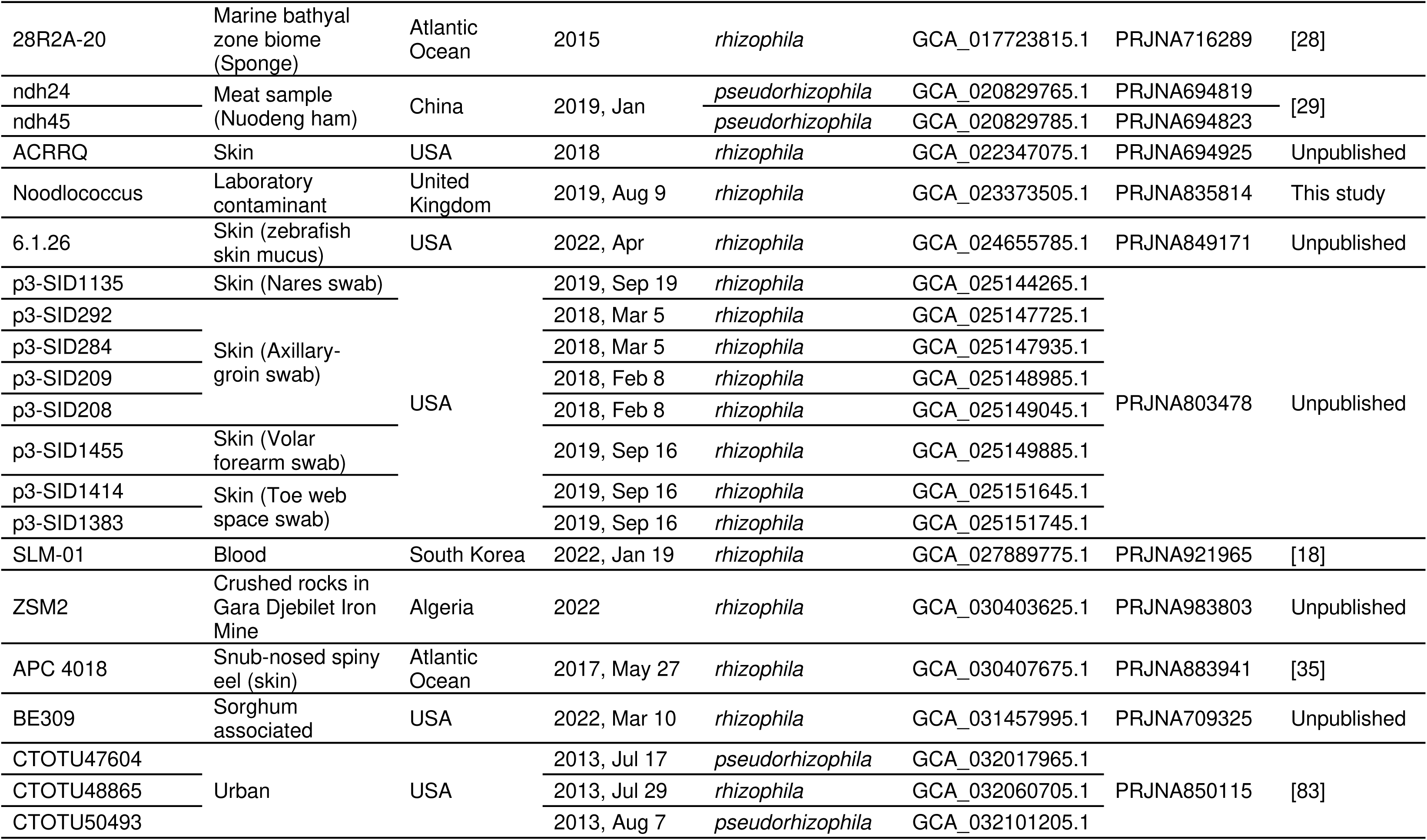
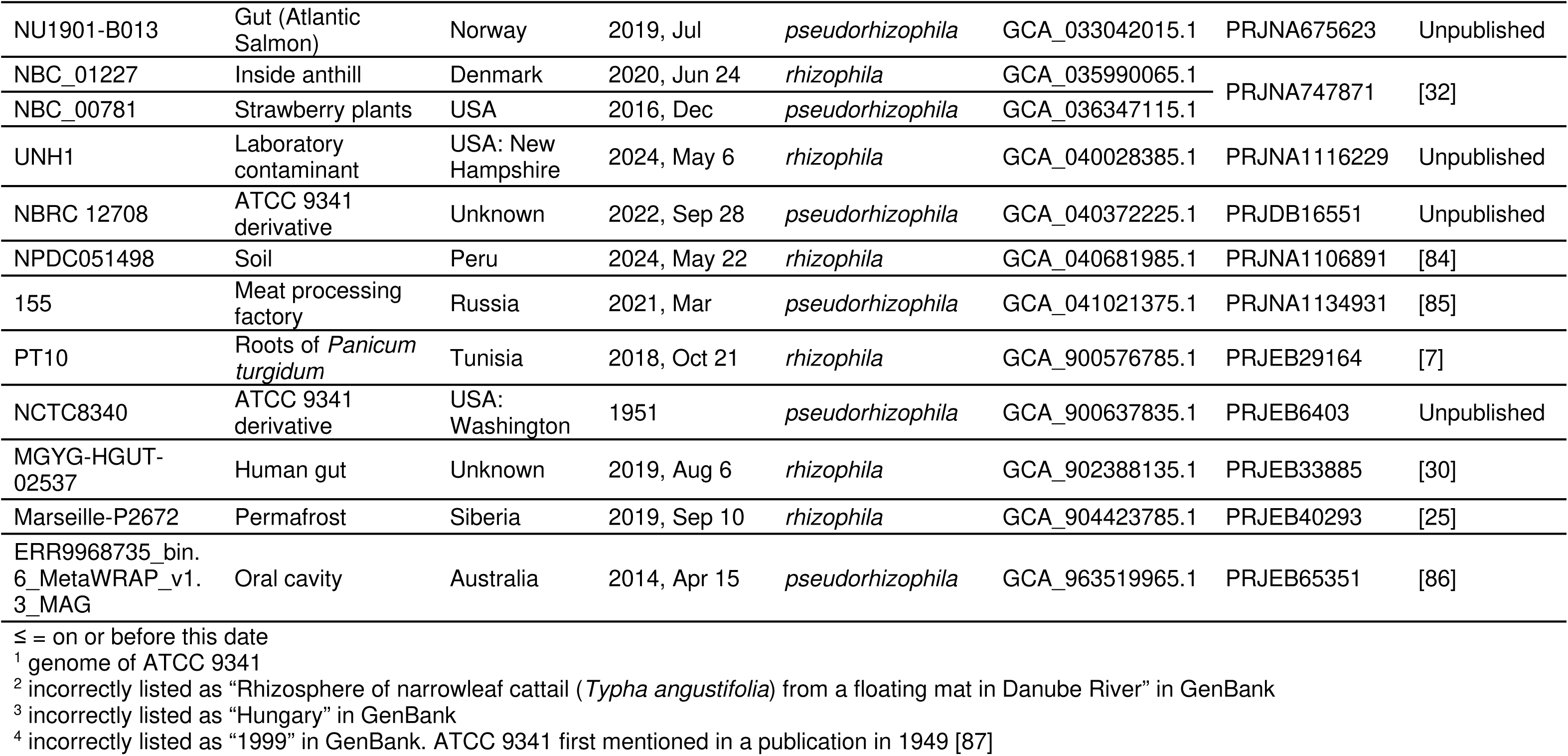
*K. rhizophila* genomes in NCBI database (last search 21st September 2024)

### Distinction of two monophyletic clades in *Kocuria rhizophila*

Genomes that have been labelled *K. rhizophila* clearly separated into two monophyletic clades in the genus-wide core genome phylogeny (Fig. S3). This topology remained when a core genome phylogeny was inferred for *K. rhizophila* genomes alone (Fig. 3), as well as in the full 16S rRNA phylogeny (Fig. S5). Type strain TA68, Noodlococcus, and commonly used reference strain NBC_01227 were located together in one monophyletic clade, whilst the genomes of *K. rhizophila* available from major culture collections (NCTC 8340 and NBRC 12708) and other commonly used reference strains (DC2201, FDAARGOS_302) were located in the other clade. FastANI revealed inter-clade ANI values of 89.5-90.4% and intra-clade ANI values of 96.7-100% (Fig. 3, Fig. 4). These ANI values clearly contrast with those observed for all other *Kocuria* species (Fig. S4), which suggests that the *K. rhizophila* clades constitute two distinct species. We refer to the clade containing *K. rhizophila* type strain TA68 as *Kocuria rhizophila*, and the other clade as *Kocuria pseudorhizophila*. We identified a total of 35 *K. rhizophila* and 16 *K. pseudorhizophila* genomes (Fig. 3, Table 1).

**Fig. 3.**
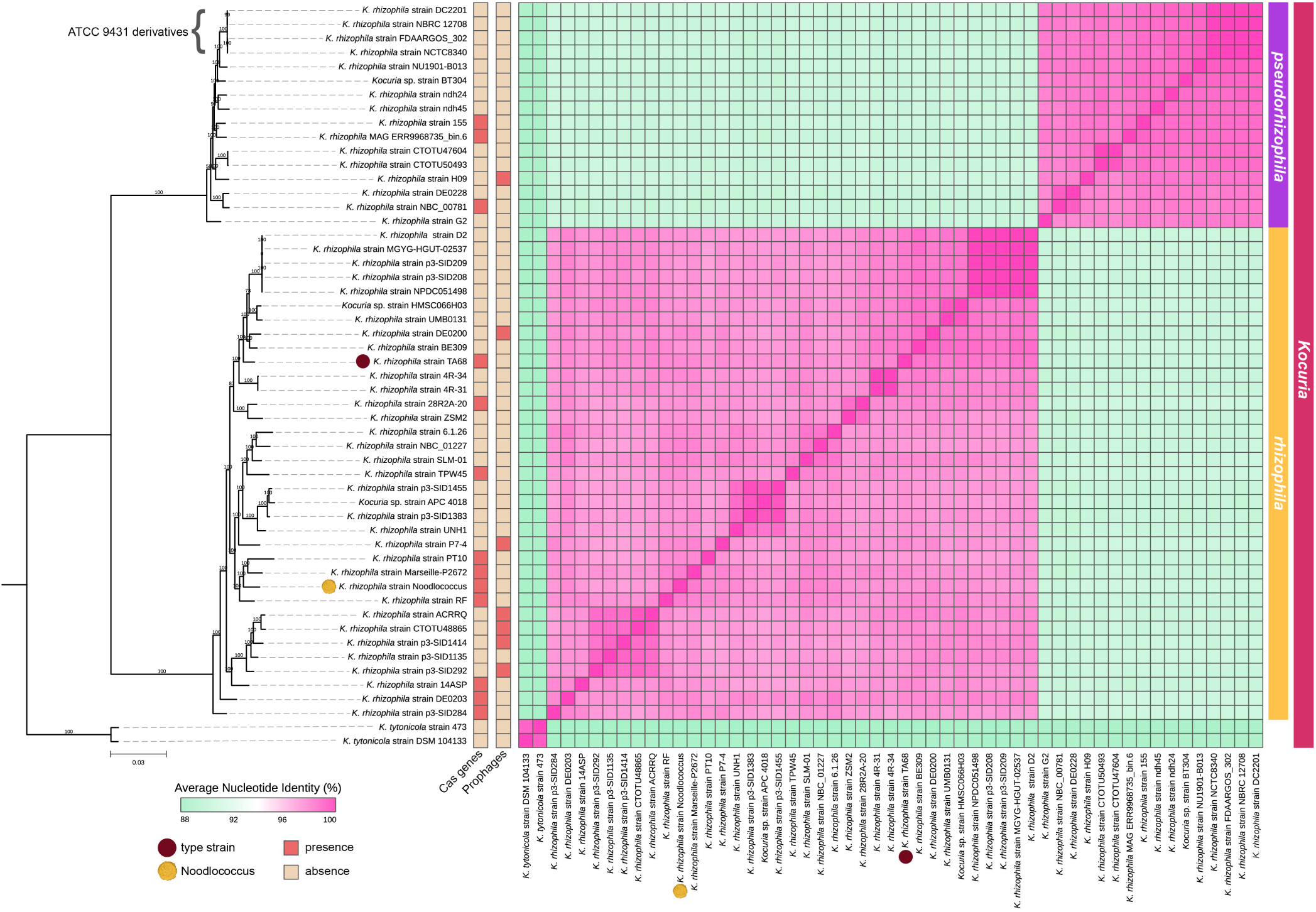
Core genome phylogeny and pairwise average nucleotide identities of *Kocuria rhizophila* genomes (*n=*51). Maximum Likelihood phylogeny was inferred with the GTR+F+I+R5 substitution model using IQTREE. Ultrafast bootstrap supports are labelled on each branch and scale bar represents substitutions per site. *Kocuria tytonicola* strains 473 and DSM 104133 were used as outgroups. Pairwise average nucleotide identities were calculated with FastANI and visualised with pheatmap. Proposed species demarcation is indicated by the group labels on the right side of the figure. Presence/absence of CRISPR (Cas genes) and prophages are shown to the right of the tree. Type strain TA68 is highlighted with a brown circle, and Noodlococcus is highlighted with a yellow circle.

**Fig. 4.**
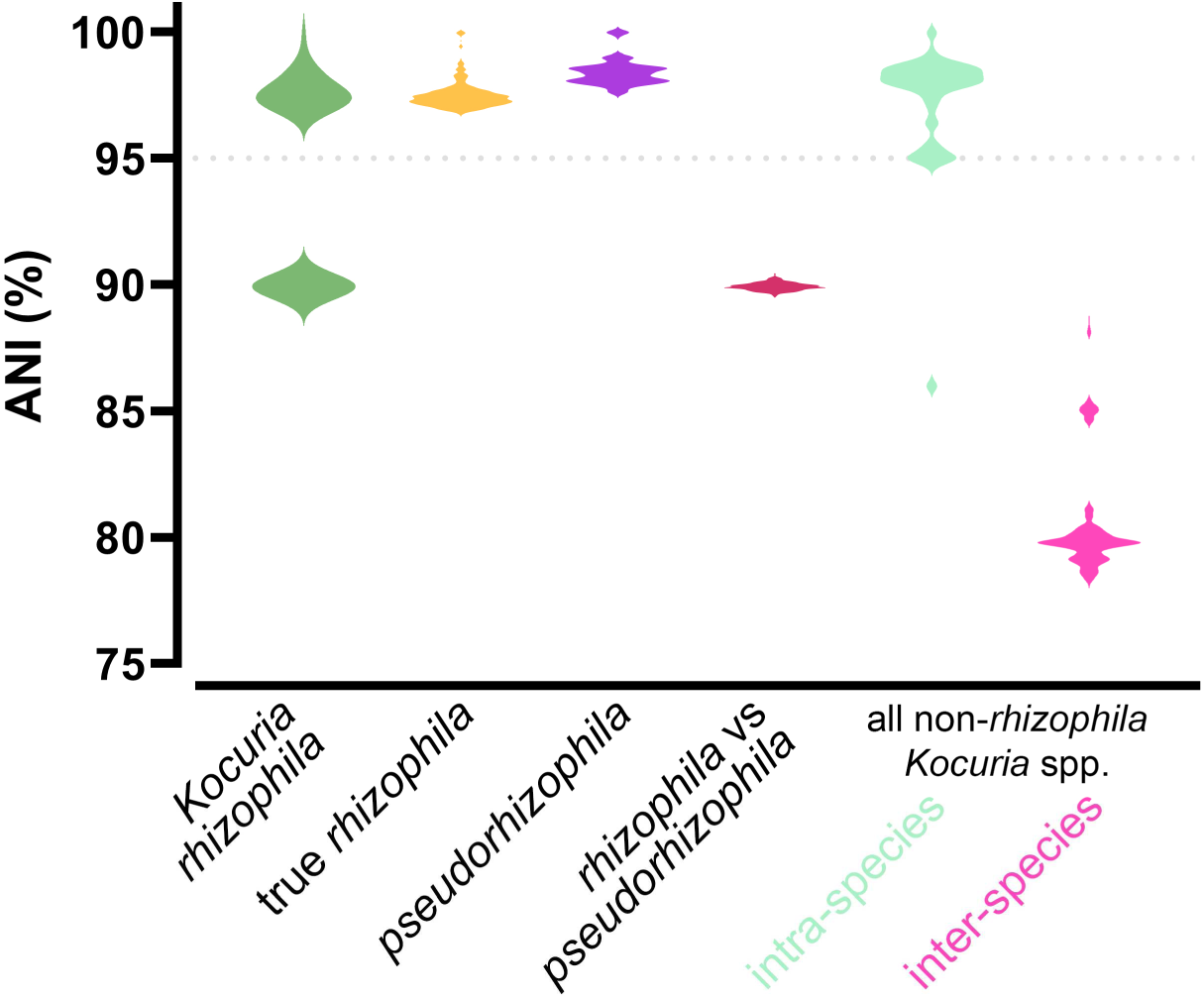
Average nucleotide identities (ANIs) between various *Kocuria* taxa. *K. rhizophila* (*n=*51), true *rhizophila* (*n=*35)*, pseudorhizophila* (*n=*16). All non-*rhizophila Kocuria* spp. only includes genomes of species groups that contained >2 genomes (*n=*141).

Within clades, we observed two clusters of genomes that exhibited near-identical ANIs. The first of these was in the *K. pseudorhizophila* clade and contained all four genomes derived from the widely used reference strain ATCC 9341 (DC2201, NBRC 12708, FDAARGOS_302, and NCTC8340), which had ANIs around 100% (Fig. 3). The second cluster included five genomes in the *K. rhizophila* clade (strains D2, MGYG-HGUT-02537, p3-SID209, p3-SID208, and NPDC051498), with ANIs ranging between 99.9-100% (Fig. 3). Interestingly, these isolates originated from a range of sources and geographical locations (Table 1, Table S5). MGYG-HGUT-02537, collated and uploaded to NCBI as part of the Unified Human Gastrointestinal Genome collection [30], was identical to and is likely a duplicate genome of strain D2 (ANI 100%), isolated from a human stool sample in India. However, NPDC051498, from the Natural Products Discovery Centre collection [84], was isolated from soil in Peru, and strains p3-SID209 and p3-SID208 were isolated from a human skin swab in the USA (Table 1, Table S5). There were nine further instances where strains from different studies shared ANIs >99% (Fig. 3, Table S5).

### Pangenome analysis further distinguishes *K. rhizophila* clades

To further interrogate the differences between the two *K. rhizophila* clades, we examined their combined pangenome. A pangenome, composed from 51 genomes from the two *K. rhizophila* clades, was constructed using Panaroo [59]. The pangenome sample consisted of 1,013 core, 711 soft core, 951 shell, and 2,275 cloud genes. Within the pangenome we identified clade-exclusive gene sets (Fig. 5). For example, *K. rhizophila* and *K. pseudorhizophila* possessed 95 and 79 clade-specific core genes, respectively. Clade-specific core genes are present in all representative genomes of one clade and never present in the other clade. Most (*n=*90/95 and *n=*63/79) clade-specific core gene clusters were unable to be assigned to clusters of orthologous genes (COG) categories, highlighting the need for further functional studies in these organisms. Notable core genes specific to the *K. pseudorhizophila* clade related to energy production and conversion (*ssuD, glcD, mauE*) and amino acid transport and metabolism (*soxE, apeB, lys2B)*. Sets of clade-specific intermediate and clade-specific rare genes were also identified (*K. rhizophila*: 326 clade-specific intermediate, 1,243 clade-specific rare; *K. pseudorhizophila*: 179 clade-specific intermediate, 623 clade-specific rare).

**Fig. 5.**
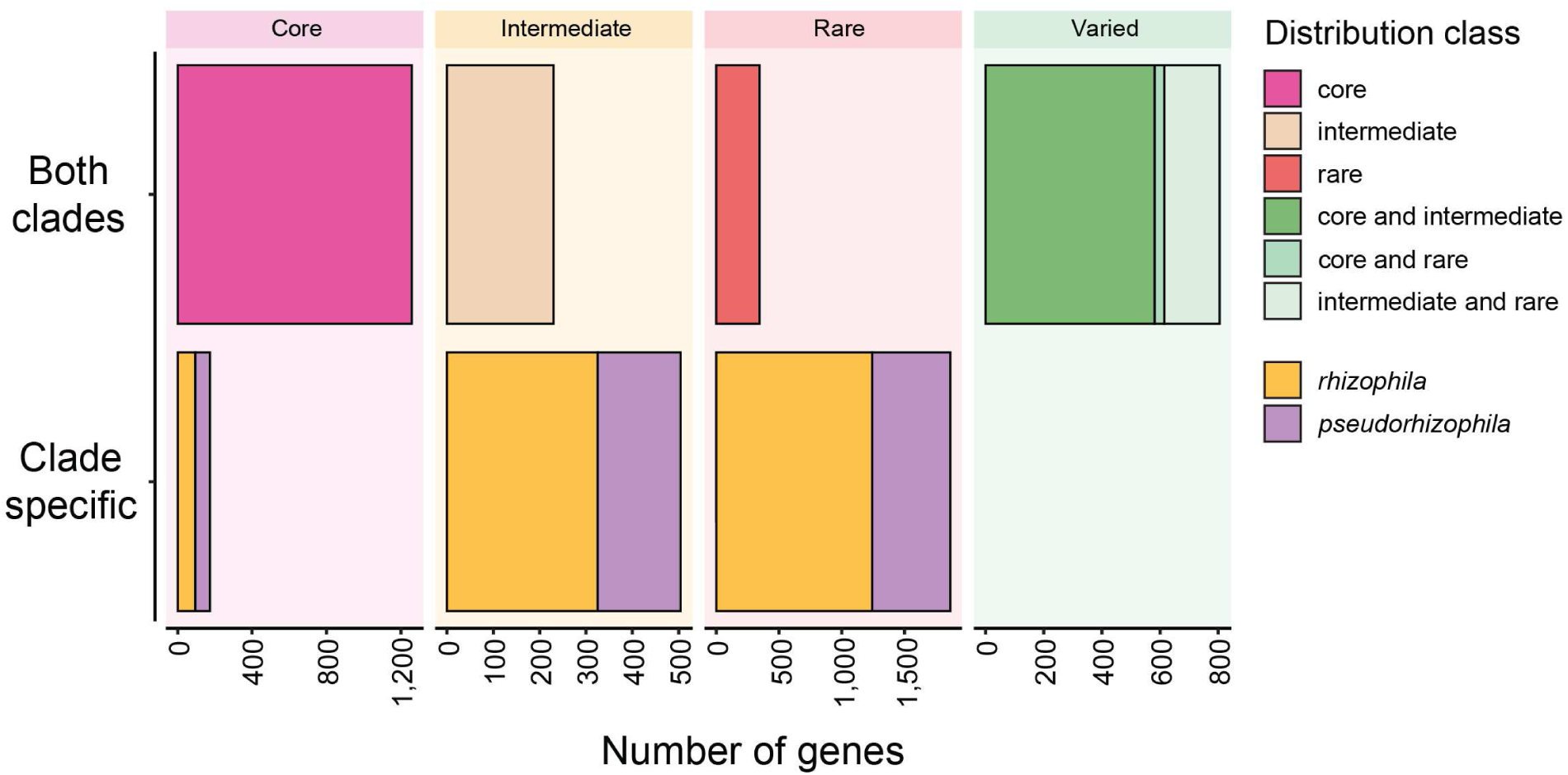
Population-structure aware pangenome of *Kocuria rhizophila*. Number of gene clusters of the *K. rhizophila* pangenome (both clades) from each distribution class. Clade-specific gene clusters exclusively present in either *rhizophila* (yellow) or *pseudorhizophila* (purple) are shown for each distribution class. Distribution class definitions taken from [68].

To examine the conservation of a characterised region of the Noodlococcus chromosome in the rest of *K. rhizophila*, we screened all 51 genomes for the presence of Cas determinants using the Noodlococcus sequence (Fig. 2c) as a reference. The Cas genes were detected in 13 genomes spread across the phylogeny (Fig. 3). The Cas gene segments in these genomes ranged from 99.4-100% identical to the region from Noodlococcus, with the exception of the segment in strain 14ASP, which was 90.6% identical. Having found no prophages in Noodlococcus, we extended our analysis to the remaining 50 genomes. Prophages were detected in six genomes in the *K. rhizophila* clade and one genome in the *K. pseudorhizophila* clade. All prophages were classified within the *Caudoviricetes* class, and their completeness scores ranged from 33.3-84.0%. Notably, three *K. rhizophila* genomes (strains DE0200, ACRRQ, and p3-SID1414) harboured the same prophage, sharing over 99% nucleotide identity. All seven prophage sequences exhibited high nucleotide identities (86.2% to 97.5%) to bacteriophage sequences identified in an urban environment metagenomic study — the same study from which phage genomes that matched Noodlococcus CRISPR spacer sequences were derived. None of the seven prophage-containing genomes contained Cas genes (Fig. 3).

## Discussion

Our serendipitous isolation and subsequent characterisation of laboratory contaminant Noodlococcus led to the first large-scale genomic assessment of the *Kocuria* genus. Despite its ubiquity in nature, *Kocuria* is obscure relative to the medically-relevant bacterial genera that account for the vast majority of genomes in public sequence databases [88]. Noodlococcus was determined to be a strain of *K. rhizophila*, and this prominent *Kocuria* species became the focus of our analyses.

The *K. rhizophila* type strain, TA68, was isolated in the 1990s [3] and its draft genome sequence is available [81]. At the time of our study, there were 50 draft and just 9 complete *K. rhizophila* genomes in NCBI, including Noodlococcus. We have shown here that, of the complete genomes available, only 28R2A-20, NBC_01227, UNH1, and Noodlococcus represent the true *K. rhizophila* clade that includes TA68 (Fig. 3). The Noodlococcus genome is the first of these to be characterised and described. Our annotations of genomic features associated with replication, carotenoid synthesis and defence are the first for this species (Fig. 2). By examining CRISPR spacers, we have identified putative *Kocuria* phages (Table S4). These phages have previously been predicted to be lytic, but were also predicted to be *Arthrobacter* phage using vHULK [76]. Given that vHULK was modelled from a relatively small number of genera that did not include *Kocuria* [89], we expect that the identification of these phage sequences in the Noodlococcus spacer region provides stronger evidence that *Kocuria* is their natural host. We also identified several putative lysogenic phages in *K. rhizophila* genomes. We used the Noodlococcus genome to explore IS, which are arguably the simplest self-mobile genetic elements. This led to the identification of four novel elements, IS*Krh4-7* (Fig. 2), which are the first examined in *K. rhizophila*, as the first three *Kocuria* IS in ISFinder (IS*Krh1-3*; all IS*481* family) were found in DC2201, which is in the *K. pseudorhizophila* clade (Fig. 3). IS*Krh4-7* therefore expand the number of IS families recognised in *Kocuria* to include IS*30* and IS*110*, the latter of which has recently been shown to have unique transposition properties and biotechnological potential [77].

Taking a broad view of the genus, we generated the largest *Kocuria* phylogeny created to date, which supported species assignments and placed some previously unidentified genomes with their closest relatives. We found that *K. rhizophila* is not a monophyletic species, and is made up of two distinct clades, which we refer to as *K. rhizophila* and *K. pseudorhizophila*. Our analyses strongly suggest that these should be classified as two different species. This was indicated by both our core genome and full 16S rRNA phylogenies. ANI comparisons also showed a clear distinction between genomes in the *K. rhizophila* and *K. pseudorhizophila* clades, with intra-clade ANI values of >96.7% and >97.5%, respectively. The universal species boundary of 95% ANI has been disputed, with many species exhibiting intra-species ANI values below this [90]. However, we found that for *Kocuria*, non-*rhizophila* intra-species values were above 94.5%, with all inter-species ANI values <90%. Thus, our analyses support a species boundary of around 95% ANI within the *Kocuria* genus. Splitting the two *K. rhizophila* clades into separate species would maintain taxonomic consistency with the rest of the *Kocuria* genus, with intra-clade ANI values >96% and inter-clade values <90.5%. The identification of distinct clade-dependent gene sets may reflect exposure to independent gene pools due to habitation of differing ecological niches. Conceivably, these two *K. rhizophila* clades may be undergoing different evolutionary trajectories, adding weight to the argument that they should be considered as two separate species.

*K. rhizophila* is commonly used as a reference strain in industrial applications, including sterility and antimicrobial susceptibility tests. However, there are multiple reference strains available to purchase, which we have shown are genetically distinct. Type strain TA68, from the true *K. rhizophila* clade (Fig. 3), is available in various culture collections under the names ATCC BAA-50, DSM 11926, IFO 16319, CCM 4950, and NBRC 16319. Genomes for the other widely used reference strain ATCC 9341 (including pseudonyms and derivatives NBRC 103217, NBRC 12708, NCTC 8340, DSM 348, DC2201, and FDAARGOS_302) clustered in the *K. pseudorhizophila* clade (Fig. 3). ATCC 9341 is widely available in commercial products for QC testing, such as *K. rhizophila* Culti-Loops™ from Thermo Fisher Diagnostics (catalogue #R4604075). This strain was originally deposited to ATCC as *Sarcina lutea*, before being reclassified as *Micrococcus luteus*. In 2003, it was reclassified again to *K. rhizophila* after DNA hybridisation experiments indicated it was more closely related to TA68 than to the *M. luteus* type strain [36]. This reclassification may have provided the basis for subsequent mislabelling of various strains. In 2008, ATCC 9341 derivative DC2201 was the first complete *K. rhizophila* genome published [37], and its use as a species reference has led to strains in the *K. pseudorhizophila* clade being misclassified as *K. rhizophila*. Adding to the confusion, the genome for ATCC 9341 (FDAARGOS_302), was uploaded to NCBI in 2018 with identical metadata to TA68 (Table 1), which is clearly a different strain. Strains from the two clades have been unknowingly used interchangeably in multiple studies. For example, a recent study of antimicrobial resistance in *Kocuria* spp. used ATCC 9341 (*K. pseudorhizophila* clade) during laboratory experiments, but used the genome of strain 4R-31 (*K. rhizophila* clade) during bioinformatic analyses [85]. Such misrepresentation could lead to conflicting results and inaccurate conclusions. This will continue to be an issue in both research and industrial applications whilst culture collections and QC products list strains such as ATCC 9341 as *K. rhizophila*, with no indication that they are genetically distinct from true *K. rhizophila*. Standardising the use of *K. rhizophila* reference strains and genomes is therefore essential.

ATCC 9341 was first mentioned (as *S. lutea* PCI 1001) in a 1949 publication by Randall and colleagues that described its use in antibiotic sensitivity testing [87]. A 1954 publication confirms that this strain was originally isolated by W. A. Randall [91]. As our study strongly indicates the need for the separation of *K. rhizophila* into two species, we suggest that an appropriate species name for the *K. pseudorhizophila* clade that includes ATCC 9341 would be *Kocuria randallii* (ran.dal’li.i. N.L. gen. masc. n. *randallii*, of Randall, named in honour of Dr William A. Randall Sr. for contributions to antibiotic assay development and regulatory microbiology).

Given the sporadic and widespread derivations of genomes captured in this relatively small dataset, we were surprised to find that several *K. rhizophila* isolates from disparate sources shared ANI values >99% (Fig. 3, Table S5). This suggests that environmental *K. rhizophila* clones can have extensive geographic distributions. These might be explained by associations with human and animal migration, or by the detection of *Kocuria* in air and clouds [92, 93], where cells would be subjected to global air currents that could contribute to their dissemination. While this is an intriguing possibility, stronger evidence supporting the distribution of individual clones will be required before it can be considered more seriously.

This study, prompted by our characterisation of a laboratory contaminant, exemplifies how chance findings, common yet often undervalued in biological research, can yield novel insights. The discovery of Noodlococcus led to the creation of “Contamination Club” (ContamClub), a social media initiative that has been a useful vehicle for professional and public science engagement. We hope that ContamClub and the story of Noodlococcus will continue to promote investigation of the unusual and understudied, particularly in the genomics era where relatively low-cost comparative studies can yield significant findings.

## Conclusions

We have found that *K. rhizophila* is not a monotypic species, but is comprised of two clades with distinct ANIs and pangenomic profiles. Distinguishing these clades has important implications for the use of *K. rhizophila* strains as controls in research and industry. The complete genome sequence of laboratory contaminant Noodlococcus has been the basis for our description of previously uncharacterised features of the *K. rhizophila* genome. The Noodlococcus genome and our large-scale genomic evaluation can serve as a baseline for future studies into the distribution, diversity, and evolution of this ubiquitous species.

## Acknowledgements

We thank all followers of ContamClub for their encouragement, inspiration, and support. Thanks to Paul Stanley for their assistance with electron microscopy.

## Author Contributions

R.A.M. serendipitously isolated Noodlococcus. The project was conceived by R.A.M., G.E.M., S.F.S.H., R.S.M., and W.V.S. Wet lab experiments were carried out by A.W., under the supervision of G.E.M., S.F.S.H., R.S.M., and R.A.M. Long-read sequencing was carried out by J.Q. Sequence annotation was performed by R.A.M. and C.W. Bioinformatic analyses were carried out by G.E.M., S.F.S.H., E.A.C, and R.A.M. Data visualisation was by G.E.M., S.F.S.H., R.A.M., E.A.C., and L.Y.S.T. The manuscript was drafted by G.E.M, S.F.S.H, and R.A.M, and was edited and revised by all authors.

