## Supplementary Figures for "The Kocurious case of Noodlococcus: genomic insights into *Kocuria rhizophila* from characterisation of a laboratory contaminant"

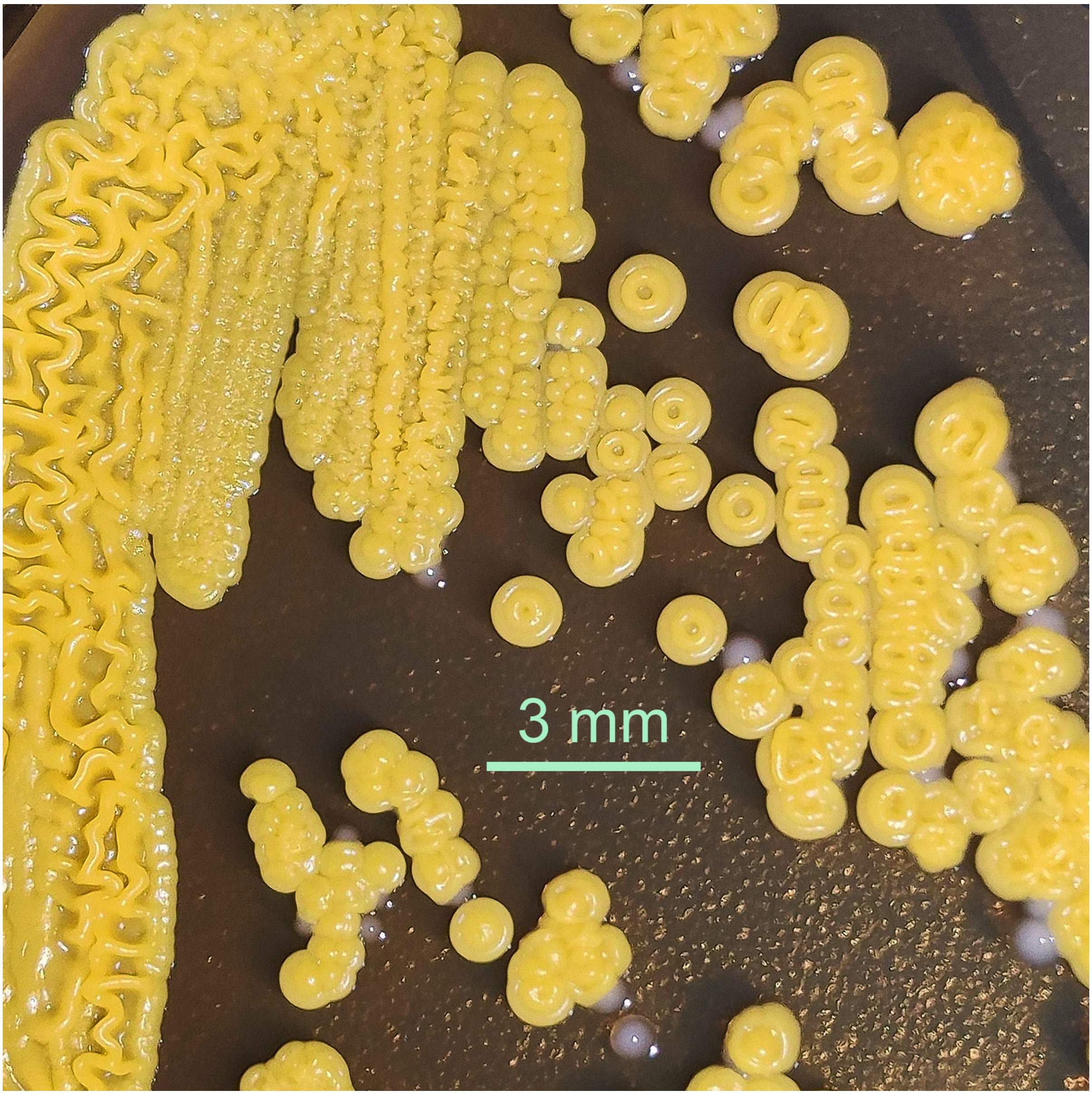


**Fig. S1. Noodlococcus colonies after 7 days of growth.** Photograph of a brain heart infusion agar plate 7 days after subsequent streaking of the original Noodlococcus colony, showing a raised central ring structure. The smaller, white colonies were *Enterococcus faecium* that had carried over from the original Noodlococcus plate.

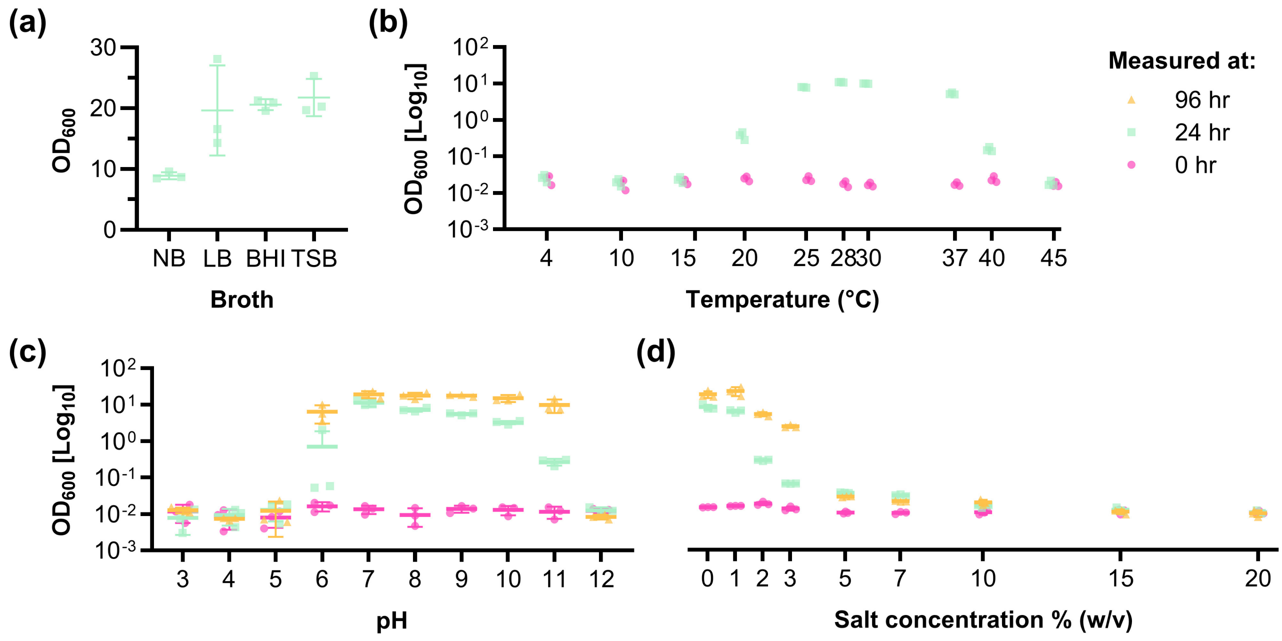


**Fig. S2. Optimal growth conditions of Noodlococcus.** Each graph shows optical density at 600 nm ($OD_{600}$) of 1 mL culture after 0, 24, or 96 hours (hr). Growth was measured **(a)** in different media (NB = nutrient broth, LB = lysogeny broth, BHI = brain heart infusion, TSB = tryptic soy broth), **(b)** at different temperatures, **(c)** in different pH conditions, and **(d)** in different weight/volume (w/v) salt concentrations. Each point represents a biological replicate as an average of three technical replicates. Error bars shown mean ± standard deviation.

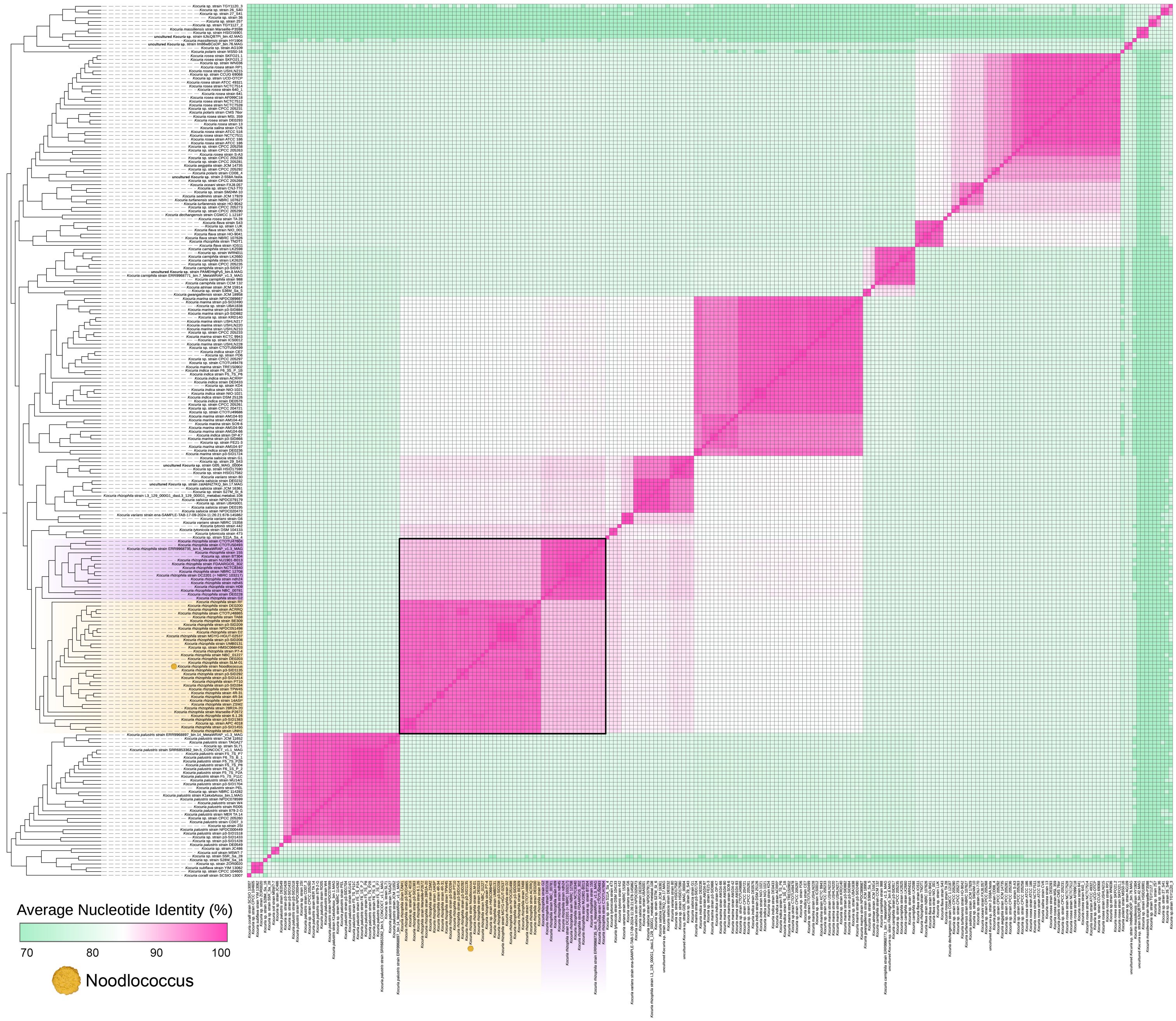


**Fig. S3. Core genome cladogram and pairwise average nucleotide identities of *Kocuria* genomes (*n*=230).** Cladogram showing the phylogenetic topology of *Kocuria* genomes, constructed from a maximum likelihood phylogeny. Pairwise average nucleotide identities (ANIs) were calculated with FastANI and visualised with pheatmap. Black box highlights the ANI values of *Kocuria rhizophila* genomes. True *K. rhizophila* genomes are highlighted in yellow, and *K. pseudorhizophila* genomes are highlighted in purple. Noodlococcus is highlighted with a yellow circle.

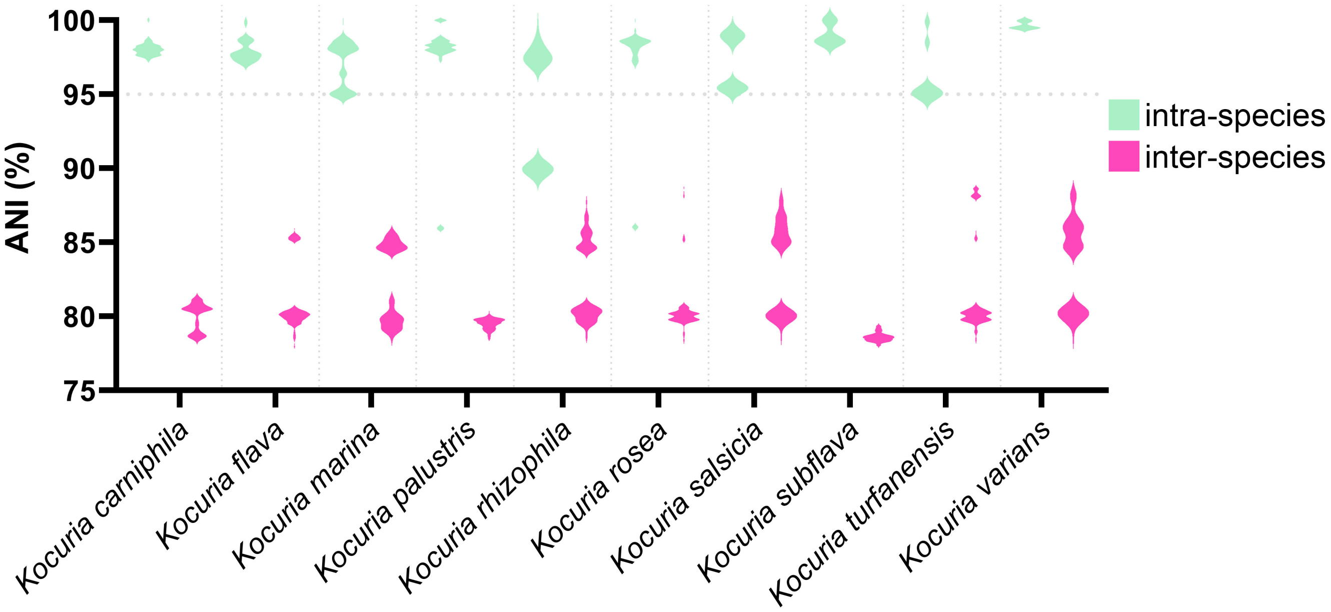


**Fig. S4. Intra-species and inter-species average nucleotide identities (ANIs) for species of *Kocuria*.** Only *Kocuria* spp. with >2 genomes were included. *K. carniphila* (*n*=10), *K. flava* (*n*=7), *K. marina* (*n*=42), *K. palustris* (*n*=28), *K. rhizophila* (*n*=51), *K. rosea* (*n*=28), *K. salsicia* (*n*=15), *K. subflava* (*n*=3), *K. turfanensis* (*n*=5), *K. varians* (*n*=3).

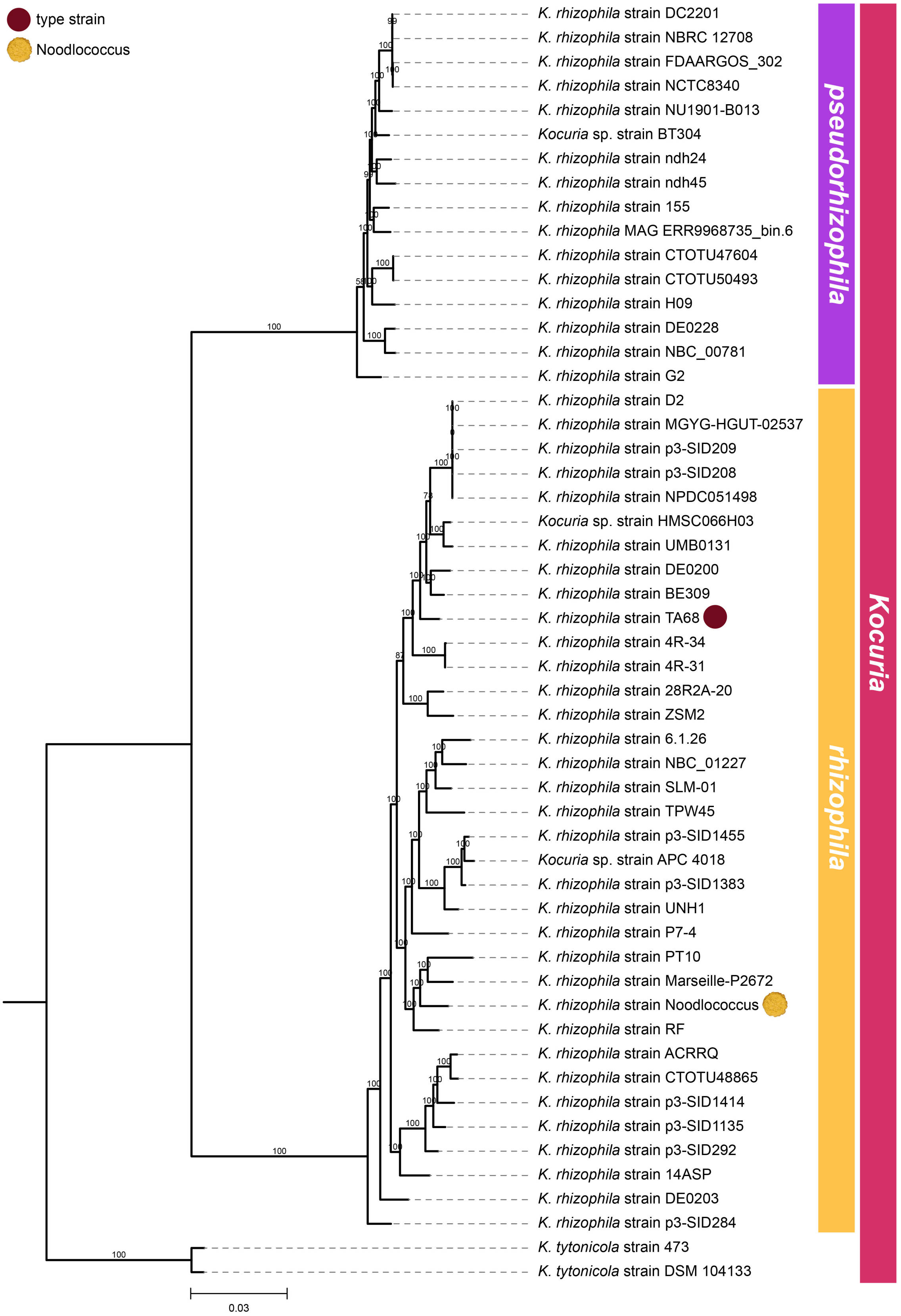


**Fig. S5. Full 16S ribosomal RNA phylogeny for *Kocuria rhizophila* genomes (*n*=51).** Ultrafast bootstrap support values are labelled on each branch and scale bar represents substitutions per site. *Kocuria tytonicola* strains 473 and DSM 104133 were used as outgroups. Proposed species demarcation is indicated by the group labels on the right side of the figure. Type strain TA68 is highlighted with a brown circle, and Noodlococcus is highlighted with a yellow circle.
